# The spermidine acetyltransferase SpeG regulates transcription of the small RNA RprA

**DOI:** 10.1101/462937

**Authors:** Linda I. Hu, Ekaterina V. Filippova, Joseph Dang, Sergii Pshenychnyi, Jiapeng Ruan, Olga Kiryukhina, Wayne F. Anderson, Misty L. Kuhn, Alan J. Wolfe

**Affiliations:** Department of Microbiology and Immunology, Loyola University Chicago, Health Sciences Division, Stritch School of Medicine, Maywood, IL 60153, USA; Center for Structural Genomics of Infectious Diseases, Northwestern University Feinberg School of Medicine, Department of Biochemistry and Molecular Genetics, Chicago, IL 60611, USA; Department of Chemistry and Biochemistry, San Francisco State University, San Francisco, CA 94132, USA; Recombinant Protein Production Core at Chemistry of Life Processes Institute, Northwestern University, Chicago, IL 60208, USA; Department of Microbiology-Immunology, Feinberg School of Medicine, Northwestern University, Chicago, IL 60611, USA; Department of Biochemistry and Molecular Biology, Knapp Center for Biomedical Discovery, University of Chicago, Chicago, IL 60637, USA; Yale University School of Medicine, Department of Digestive Diseases, New Haven, CT 06510, USA

## Abstract

Spermidine *N*-acetyltransferase (SpeG) acetylates and thus neutralizes toxic polyamines. Studies indicate that SpeG plays an important role in virulence and pathogenicity of many bacteria, which have evolved SpeG-dependent strategies to control polyamine concentrations and survive in their hosts. In *Escherichia coli*, the two-component response regulator RcsB is reported to be subject to Nε-acetylation on several lysine residues, resulting in reduced DNA binding affinity and reduced transcription of the small RNA *rprA*; however, the physiological acetylation mechanism responsible for this behavior has not been fully determined. Here, we performed an acetyltransferase screen and found that SpeG inhibits *rprA* promoter activity in an acetylation-independent manner. Surface plasmon resonance analysis revealed that SpeG can physically interact with the DNA-binding carboxyl domain of RcsB. We hypothesize that SpeG interacts with the DNA-binding domain of RcsB and that this interaction might be responsible for SpeG-dependent inhibition of RcsB-dependent *rprA* transcription. This work provides a model for SpeG as a modulator of *E. coli* transcription through its ability to interact with the transcription factor RcsB. This is the first study to provide evidence that an enzyme involved in polyamine metabolism can influence the function of the global regulator RcsB, which integrates information concerning envelope stresses and central metabolic status to regulate diverse behaviors.

## Introduction

SpeG, a member of the Gcn5-related *N-*acetyltransferase (GNAT) family, is a bacterial spermidine *N*-acetyltransferase that acetylates spermidine and spermine. These polyamines are toxic to bacteria at high concentrations and acetylation neutralizes this toxicity [1, 2]. Studies indicate that SpeG plays an important role in virulence and pathogenicity of many bacteria, which have evolved SpeG-dependent strategies to control polyamine concentrations and survive in their hosts [3-6]. Kinetic and structural analyses have demonstrated that SpeG from both *Escherichia coli* and *Vibrio cholerae* can acetylate spermidine [7-9]. These studies also showed that SpeG from *V. cholerae* is an allosteric protein; when spermidine binds to its allosteric site, SpeG exhibits a symmetric closed dodecameric structure [7, 9]. Finally, in the absence of spermidine binding, *V. cholerae* SpeG can adopt a unique asymmetric dodecameric structure with an open conformational state [10].

During the course of this study, we found that SpeG also regulates the small RNA *rprA*, whose transcription strictly requires the phosphorylated isoform of the two-component response regulator RcsB [11, 12]. The canonical two-component signal transduction system is composed of two proteins. The first is a sensor kinase that detects a signal and, in response, autophosphorylates a conserved histidine residue using ATP as the phosphoryl donor. The second is a response regulator that autophosphorylates a conserved aspartate residue using the phosphorylated sensor kinase as the phosphoryl donor [for reviews, see [13-15]]. A more complex variant of the basic two-component system is the phosphorelay, such as the Rcs phosphorelay, which consists of five proteins (RcsC, RcsD, RcsF, IgaA, and RcsB). The first four proteins are involved in controlling the phosphorylation status of the response regulator RcsB in response to diverse extracytoplasmic stimuli. The phosphorylation status of RcsB is set by an ATP-dependent protein-protein interaction chain whose core consists of the cytoplasmic membrane-associated sensor kinase/phosphatase RcsC and its cognate histidine phosphotransferase RcsD [16]. The inner membrane protein IgaA favors RcsC phosphatase activity and thus dephosphorylation of RcsB. Relocation of the outer membrane lipoprotein RcsF to the periplasm favors RcsC kinase activity and thus phosphorylation of RcsB. This occurs when RcsF interacts with the C-terminal periplasmic domain of IgaA. Together, RcsF and IgaA regulate the activities of the Rcs phosphorelay components [17-24]. RcsB also can become phosphorylated in response to central metabolic changes via the central metabolite acetyl phosphate [25]. Both mechanisms (RcsC-dependent and acetyl phosphate-dependent) regulate the phosphorylation status of RcsB and thus both control RcsB-dependent processes, such as desiccation, flagellar biogenesis, capsule biosynthesis, and cell division [16, 25-28].

The Rcs phosphorelay is unusual, as the response regulator RcsB can form both a homodimer and a variety of heterodimers. The homodimer activates transcription of *rprA* [11, 12, 29], which encodes the small RNA regulator of the stationary phase sigma factor RpoS, and represses transcription of *flhDC*, which encodes the master regulator of the flagellar regulon [25, 30, 31]. To activate synthesis of the capsular exopolysacchararide colanic acid, RcsB forms a complex with a partner transcription regulator, RcsA, stabilizing the interaction between RcsB and a specific DNA binding site, the “RcsAB box” [32, 33]. RcsB also can form protein-protein complexes with other partner transcription factors, including GadE, RmpA, MatA, BglJ, and RflM; there is also evidence to suggest an interaction with PhoP [34-39]. Because these protein-protein complexes form in response to a variety of conditions, the Rcs system can mediate diverse responses that contribute to biofilm formation, virulence, motility and antibiotic resistance in pathogens [26-28, 34-36].

Biochemical and mass spectrometry analyses indicate that RcsB can become N^ε^-lysine acetylated on multiple residues [29, 40-42]. Two mechanisms for N^ε^-lysine acetylation have been reported. One mechanism involves the direct donation of the acetyl group from acetyl phosphate to a deprotonated lysine ε-amino group [41, 43]. The other mechanism is enzymatic, relying on a lysine acetyltransferase (KAT) to catalyze donation of the acetyl group from acetyl-coenzyme A (acCoA) to the ε-amino group of a lysine residue [44]. All known bacterial KATs are members of the large family of GNATs [29, 40, 44-47].

One of our previous studies suggested that acetylation of RcsB diminished its ability to activate *rprA* transcription in *E. coli* [29]. In an effort to identify a KAT that might be responsible, we first screened 21 known *E. coli* genes that encode or are predicted to encode GNATs, seeking those that inhibited *rprA* transcription. This screen revealed that SpeG could inhibit *rprA* activity; however, we obtained no evidence that SpeG functions as a RcsB lysine N^ε^-acetyltransferase. Instead, we report here that SpeG can interact with RcsB through the latter’s DNA binding domain. Our findings represent the first evidence that the metabolic enzyme SpeG can affect transcription by interacting with the response regulator RcsB.

## Results

### SpeG regulates *rprA* promoter activity

While the GNAT YfiQ (also known as Pka and PatZ) can acetylate RcsB *in vitro* [29, 40], the *yfiQ* mutant does not affect RcsB acetylation [29]. Therefore, we suspected another GNAT was responsible for RcsB acetylation and proceeded to test a series of 21 known or putative GNATs. We overexpressed these GNATs and measured their effect on P*rprA-lacZ*, a transcriptional fusion of the RcsB-dependent *rprA* promoter (P*rprA*) and the *lacZ* gene, which we had integrated as a single copy into the chromosome of BW25113 to generate our reference strain AJW3759 (**Table 1**) [12]. From this preliminary screen, we identified SpeG as an inhibitor of P*rprA* activity. When SpeG was overexpressed from a plasmid in the reference strain, P*rprA* activity was reduced compared to the vector control during late exponential growth and during the transition into early stationary phase (OD > 1.0, **Fig 1A**, linear regression analysis t=-2.553, p=0.01472). When *speG* was deleted, P*rprA* activity increased in the isogenic *speG* mutant compared to its wild-type parent (**Fig 1B**, linear regression analysis t=7.750, p= 8.65E-12). Based on these results, we conclude that SpeG inhibits transcription from P*rprA*.

**Table 1.**
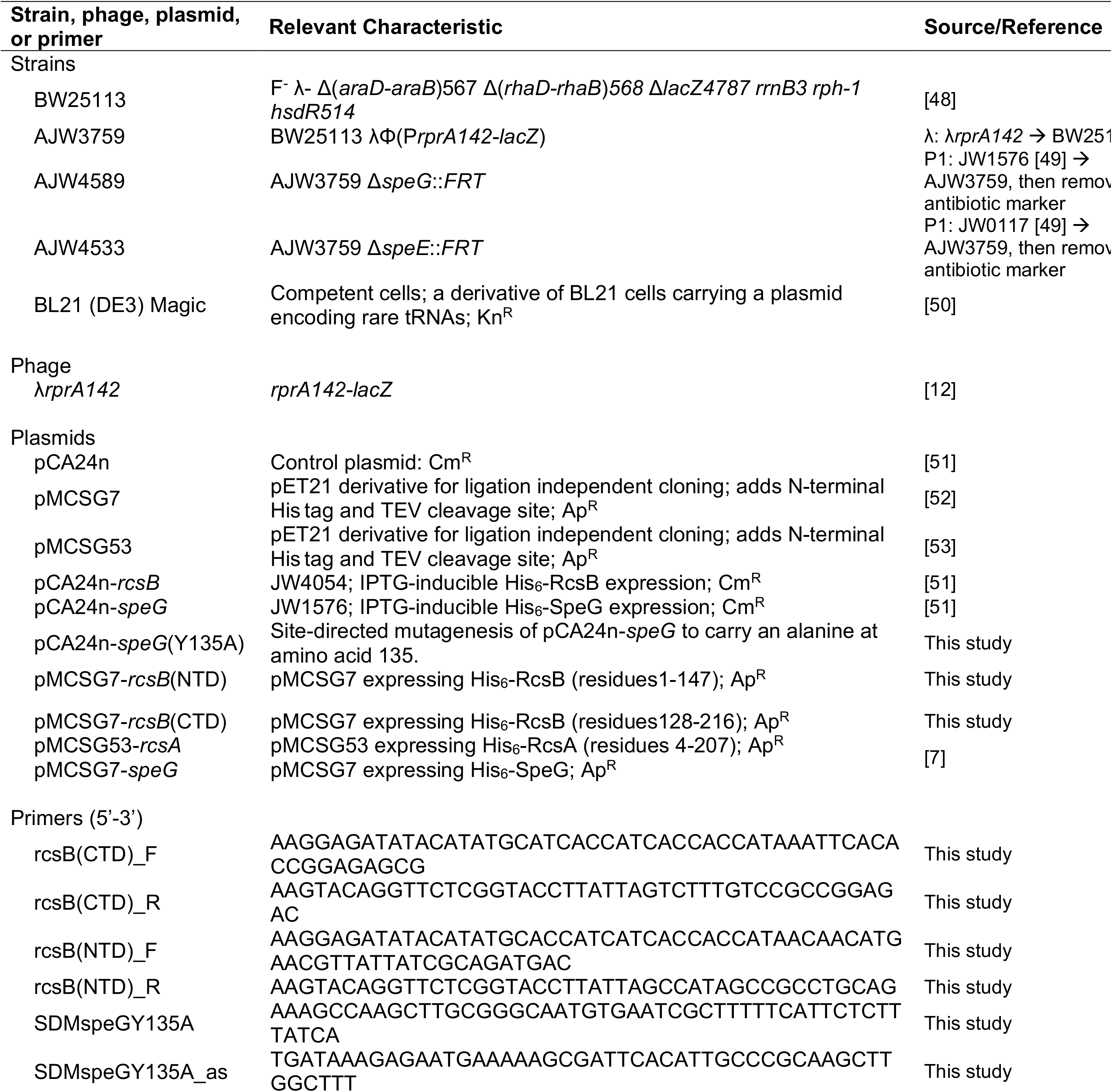
Bacterial strains, bacteriophage, plasmids, and primers used in this study.

**Fig 1.**
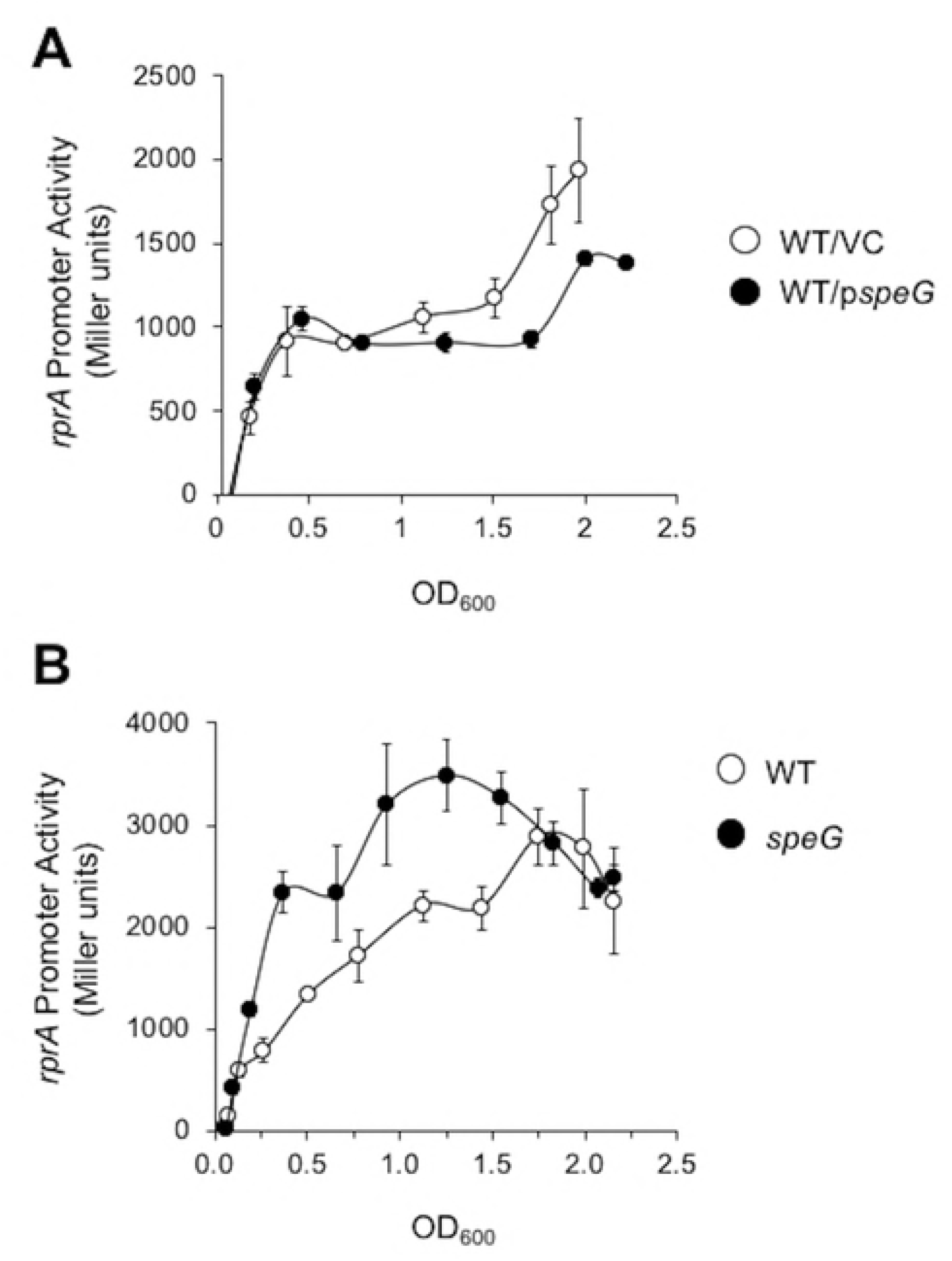
The effect of SpeG on *rprA* promoter activity. A. WT cells carrying the P*rprA*-*lacZ* fusion (AJW3759) were transformed with either a plasmid that expresses SpeG under the control of an IPTG-inducible promoter (p*speG*; pCA24n-*speG*) or the vector control (VC; pCA24n) and grown in TB7 containing 50 μM IPTG to induce SpeG expression and chloramphenicol to maintain the plasmid. Cell growth and β-galactosidase activity were assayed at various points throughout growth. The values represent average promoter activity with standard deviations of triplicate independent cultures. Linear regression analysis of the experimental group WT/p*speG* on *rprA* promoter activity versus WT/VC was statistically significant (t=-2.553, p=0.01472). B. WT (AJW3759) and isogenic *speG* (AJW4589) strains were assayed for cell growth in TB7 and β-galactosidase activity. The values represent average promoter activity with standard deviations of five independent WT and *speG* cultures. Linear regression analysis of the experimental group *speG* on *rprA* promoter activity versus WT was statistically significant (t=7.750, p= 8.65E-12).

### SpeG does not acetylate RcsB *in vitro*

Since SpeG belongs to the GNAT family of acetyltransferases known to acetylate proteins, we also tested the hypothesis that SpeG regulates *rprA* transcription by acetylating RcsB. To accomplish this, we used an *in vitro* colorimetric enzymatic assay with purified recombinant proteins. This assay measures the formation of product (CoA) indirectly via its reaction with dithionitrobenzoic acid (DTNB) to produce the thioanion product thionitrobenzoate (TNB^2-^), which is monitored spectrophotometrically at 415 nm [7, 54]. We compared the acetylation activity of SpeG toward spermidine or RcsB. As predicted, we detected SpeG acetylation activity on spermidine when acCoA was present; however, we observed no change in RcsB acetylation status in the presence of SpeG and acCoA (**Fig 2)**. This result suggests that RcsB is not a substrate for SpeG under the conditions we used to assay acetylation.

**Fig 2.**
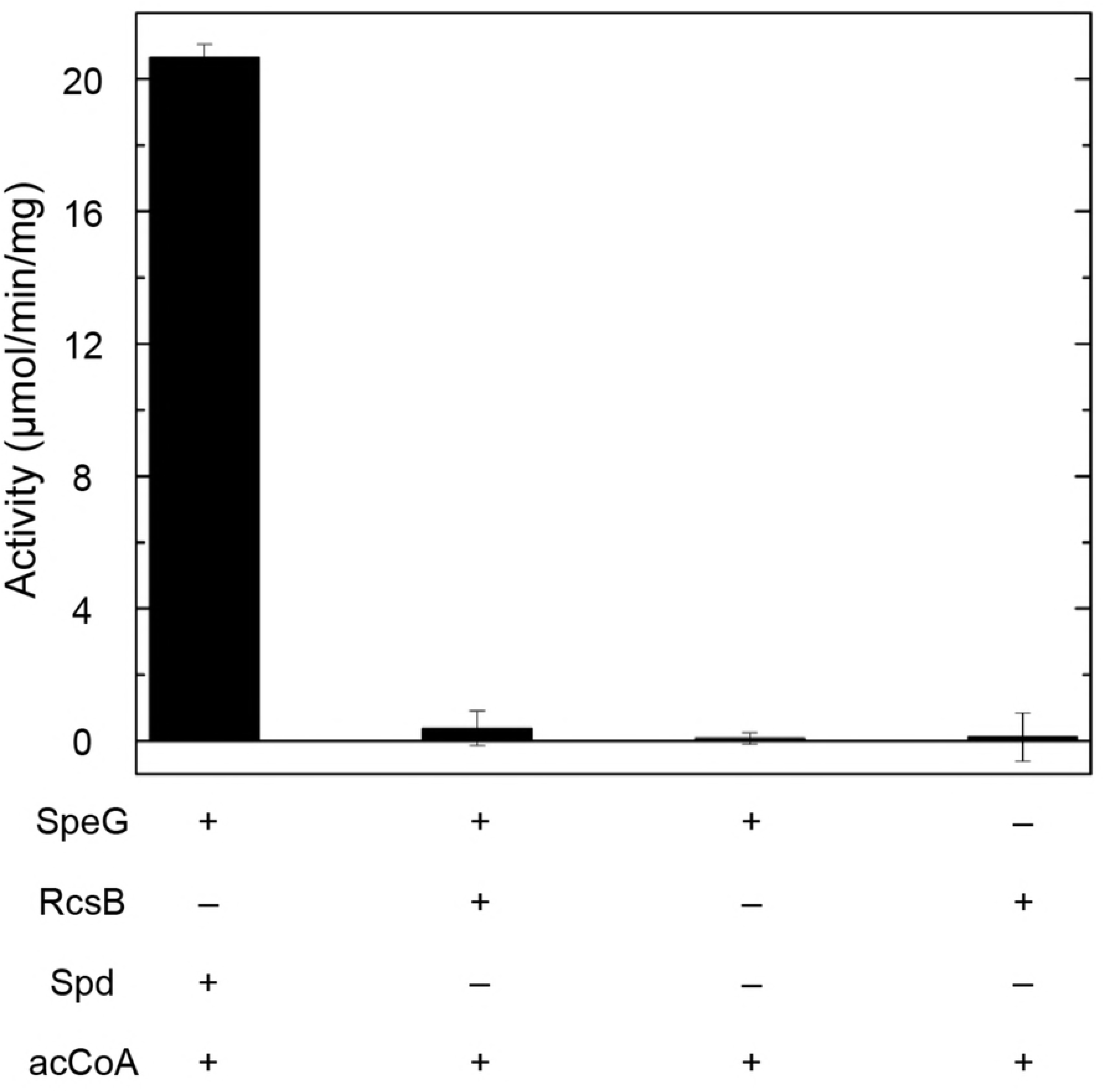
*In vitro* acetylation activity of SpeG toward RcsB or spermidine. SpeG was incubated with either spermidine or RcsB in the presence of the acetyl donor acCoA to determine if SpeG uses both spermidine and RcsB as substrates. Control reactions of possible non-enzymatic acetylation of SpeG and RcsB via acCoA were also performed. See Materials and Methods for specific reaction conditions.

### Spermidine synthase (SpeE) is not required for SpeG-dependent inhibition of *rprA* transcription

The spermidine synthase SpeE transfers a propylamine from decarboxylated S-adenosylmethionine to putrescine to form spermidine, which is both a substrate and an allosteric activator of SpeG [7]. To explore the role of SpeE/spermidine in SpeG overexpression-inhibited P*rprA* activity, we transformed a mutant that does not synthesize spermidine (*speE*) and its WT parent with either the SpeG overexpression plasmid or its vector control and monitored P*rprA* activity (**Fig 3**). SpeG overexpression resulted in reduced P*rprA* activity in both the parental strain (**Fig 3**, linear regression analysis t=-3.752, p=0.000282) and the *speE* mutant (**Fig 3**, linear regression analysis t=-3.470, p=0.000745). Furthermore, exposure of the *speE* mutant to exogenous spermidine exerted no effect on P*rprA* activity whether or not SpeG was overexpressed (**S1 Fig**). We conclude that SpeG can inhibit *PrprA* activity regardless of SpeE/spermidine status.

**Fig 3.**
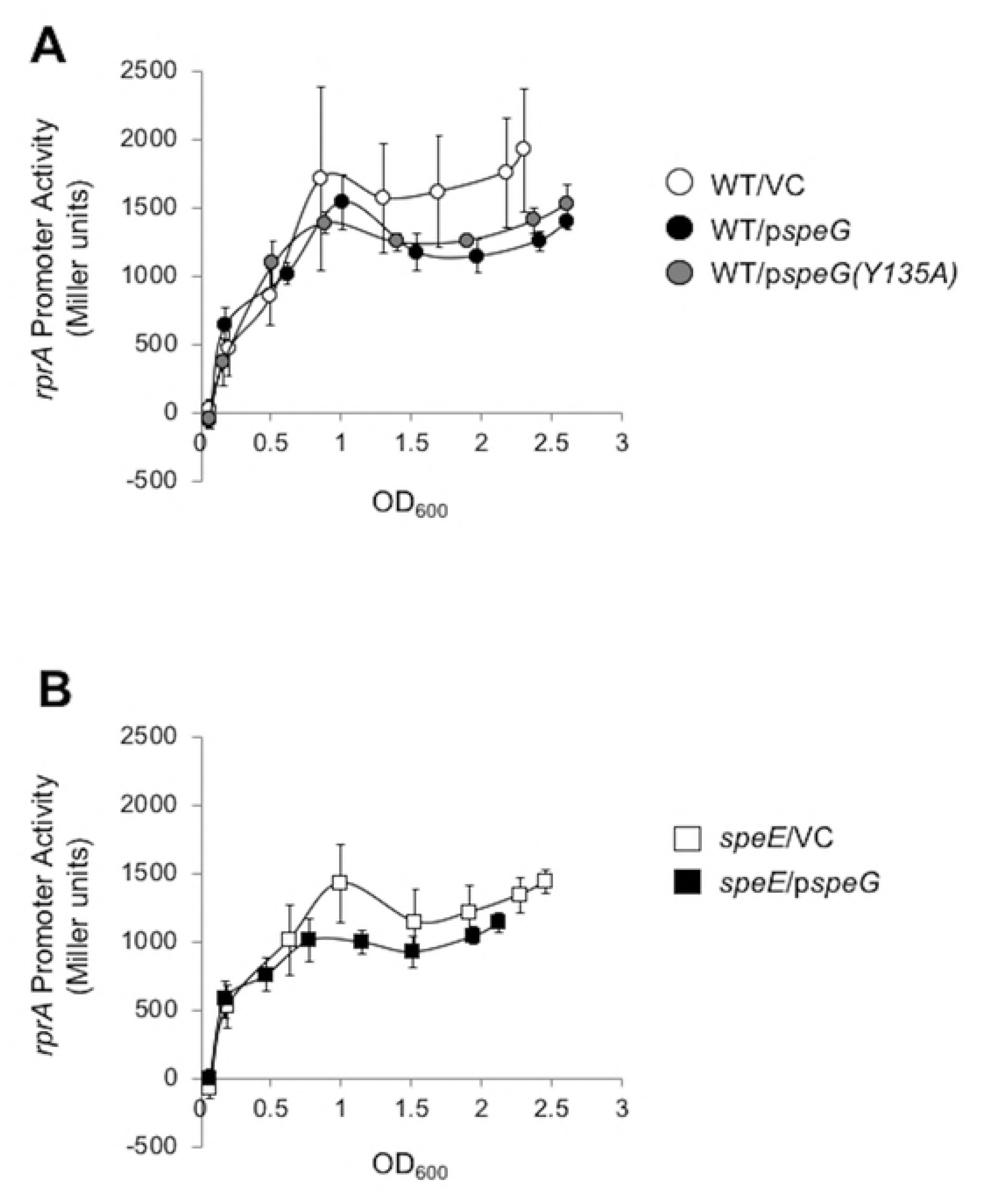
The effect of overexpressing SpeG or SpeG(Y135A) in WT cells and overexpressing SpeG in the *speE* mutant on *rprA* promoter activity. A. WT cells carrying the P*rprA*-*lacZ* fusion (AJW3759) were transformed with either p*speG* (pCA24n-*speG*), p*speG(Y135A)* (pCA24n-*speG(Y135A)*), or the VC (pCA24n) and grown in TB7 supplemented with 50 μM IPTG and chloramphenicol to maintain the plasmid. Cell growth and β-galactosidase activity were assayed. The values represent average promoter activity with standard deviations of five independent cultures. B. The isogenic *speE* mutant was transformed with either p*speG* (pCA24n-*speG*) or the VC (pCA24n) and cell growth and β-galactosidase activity were assayed as described for 3A. The values represent average promoter activity with standard deviations of five independent cultures. Linear regression comparison results on *rprA* promoter activity were significant for all experimental groups: WT/p*speG* versus WT/VC (t=-3.752, p=0.000282), WT/p*speG(Y135A)* versus WT/VC (t= -2.456, p=0.015623), and *speE*/p*speG* versus *speE*/VC (t=-3.470, p=0.000745).

### Catalytic activity of SpeG is not required for SpeG-dependent inhibition of *rprA* transcription

We next asked if SpeG overexpression-dependent inhibition of P*rprA* activity requires the spermidine acetyltransferase activity of SpeG. We therefore overexpressed SpeG Y135A, a predicted catalytically inactive SpeG variant, in the parent (AJW3759). This tyrosine (Y) residue acts as a general acid during substrate acetylation and has been shown to be critical for catalytic activity of many GNAT homologs [55-57]. We found that the SpeG Y135A mutant retained the ability to inhibit P*rprA* activity in the parent AJW3759 (**Fig 3A**, linear regression analysis t= -2.456, p=0.015623). These results are consistent with a SpeG-dependent, but spermidine acetylation-independent mechanism of inhibition in WT cells.

### SpeG binds to RcsB through its C-terminal domain

Since SpeG does not appear to acetylate RcsB and its catalytic activity is unnecessary for its ability to inhibit *rprA* transcription, we considered whether SpeG inhibits RcsB activity through a physical interaction. We used SPR to investigate whether SpeG and RcsB can form a complex. First, we immobilized SpeG onto the SPR chip and evaluated whether full-length RcsB or its N- or C-terminal domains could bind to SpeG. Both full-length RcsB (**Fig 4A**) and its C-terminal domain (**Fig 4B**) bound to immobilized SpeG in a concentration-dependent manner. In contrast, the N-terminal domain of RcsB did not (**Fig 4C**). These results suggest that RcsB binds to SpeG through its C-terminal domain. We also performed the reverse experiment, assessing whether SpeG could bind to immobilized RcsB or its domains, but we detected no signal (data not shown). Perhaps RcsB binds to the chip in a manner that prevents interaction with SpeG.

**Fig 4.**
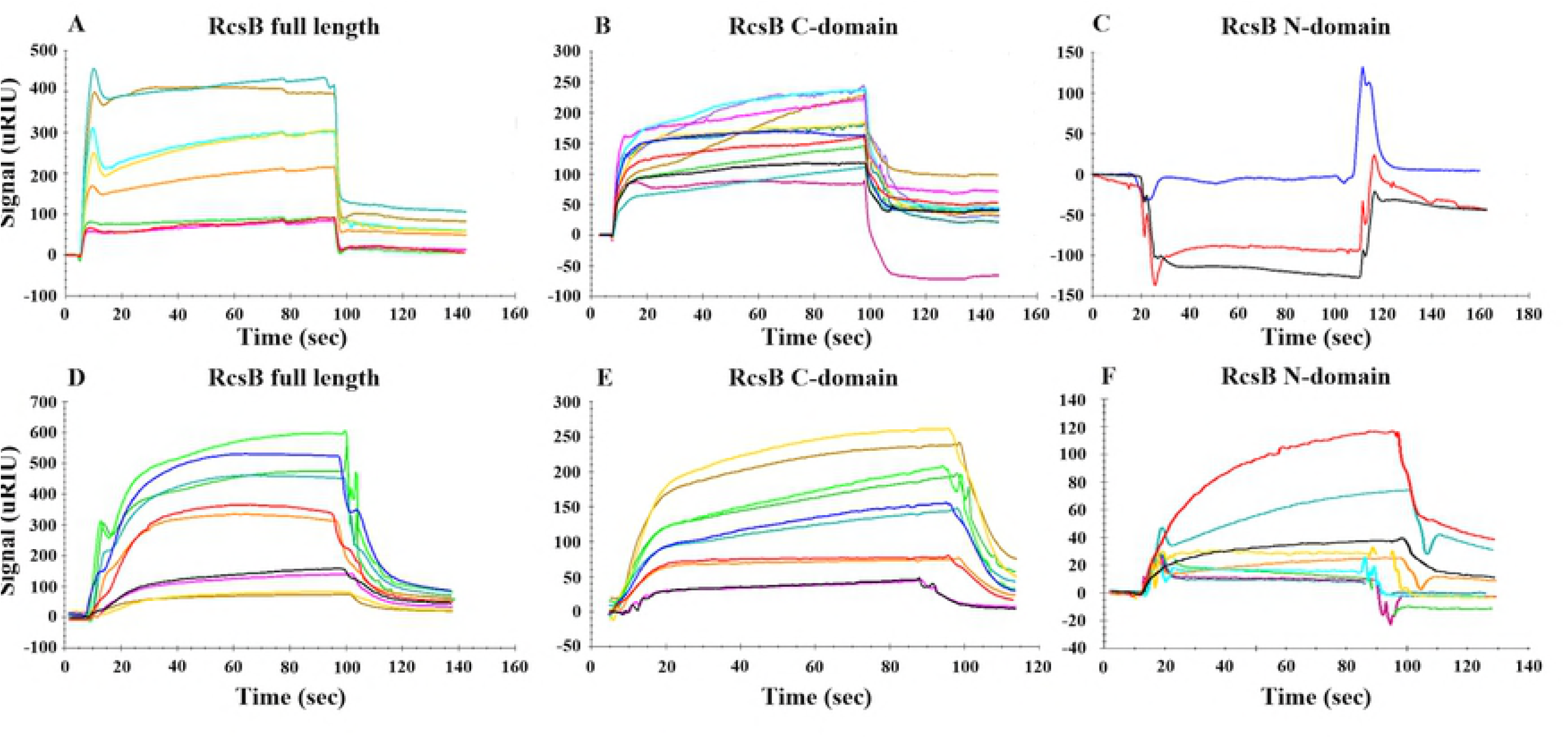
SPR analysis of the SpeG-RcsB interaction. The dose-response analysis for immobilized *E. coli* SpeG (46 μM) with increasing concentrations of full-length *E. coli* RcsB (21, 42, 53, 63 and 74 μM) as an analyte in the absence of spermidine (A) or RcsB (21, 42, 53, 63 and 74 μM) after exposure to 0.5 mM spermidine (D), RcsB C-terminal domain (45, 67, 91, 114, 136, and 159 μM) in the absence of spermidine (B) or RcsB C-terminal domain (23, 45, 91, 114, and 136 μM) after exposure to 0.5 mM spermidine (E), and RcsB N-terminal domain (59, 118, and 176 μM) in the absence of spermidine (C) or RcsB N-terminal domain (59, 118, 177, 236 and 295 μM) after exposure to 0.5 mM spermidine (F).

We calculated a K_D_ of 67 μM for the binding of SpeG to the RcsB C-terminal domain from the fit to a simple one-to-one binding model, in which one C-domain RcsB molecule interacts with one SpeG molecule (**S2A Fig**). However, we could not determine the K_D_ for the SpeG/full-length RcsB interaction, as the data did not fit either a simple binding model or other models defined in the SPR data analysis software program TraceDrawer. The lack of fitting for the SpeG/full-length RcsB interaction likely resulted from the pronounced peak at the beginning of the sensograms, which occurred especially with higher concentrations of RcsB.

We next tested the effect of spermidine on the SpeG-RcsB interaction. To accomplish this, we exposed the surface of the chip containing immobilized SpeG to spermidine and then measured the SPR signal from binding the three separate RcsB constructs (described in Materials and Methods; **Fig 4D-F**). By fitting the sensograms to these data using the one-to-one binding model, we obtained K_D_ values of 128 and 281 μM for the RcsB full-length and its C-terminal domain, respectively (**Fig S2B-C**). In contrast, we could not determine a K_D_ for the RcsB N-terminal domain due to a large chi-squared fitting value. Furthermore, we conclude that the binding of the N-terminal domain to SpeG is weak because the response signals obtained at concentrations greater than 100 μM were relatively low (**Fig 4F**). On the basis of these data and those obtained in the absence of spermidine, we propose that SpeG interacts with the C-terminal domain of RcsB in the presence or absence of spermidine and that spermidine does not prevent RcsB binding to SpeG.

### Possible SpeG inhibition of LuxR/FixJ-like transcription factors

SpeG inhibits *rprA* transcription and binds RcsB through the carboxyl terminal domain, which contains the conserved DNA binding helix-turn-helix (HTH) motif found in RcsB and other LuxR/FixJ-type proteins [16, 58]. Based on this result, it is tempting to speculate that SpeG may also bind other LuxR/FixJ family members. To identify conserved residues of the RcsB HTH motif across other LuxR/FixJ-like transcriptional regulators from *E. coli*, we used the PSI-BLAST server [59] to generate a list of DNA-binding domains from LuxR/FixJ-type family homologs and the NMR structure of the RcsB C-terminal domain from *Erwinia amylovora,* a close relative of *E. coli* [32] to visualize sequence conservation with respect to the three-dimensional structure (**Fig 5**). We also generated a phylogenetic tree using these sequences to determine which RcsB homologs had the greatest sequence similarity to its C-terminal domain and, therefore, propensity for interacting with SpeG (**S3 Fig**). We found the most conserved RcsB C-terminal domain residues across LuxR/FixJ-type homologs are S152, P153, K154, L167, V168, T169, R177, S178, K180, T181, S183, S184, Q185, K186, K187, and D198. From our analysis, the *E. coli* LuxR/FixJ-type homolog sequences of YjjQ, BglJ, YahA, YuaB, DctR, and RcsA are most similar to the RcsB C-terminal DNA-binding domain and warrant further testing. We hypothesized that SpeG might bind RcsB through these critical residues in the C-terminal domain and potentially those of other homologs.

**Fig 5.**
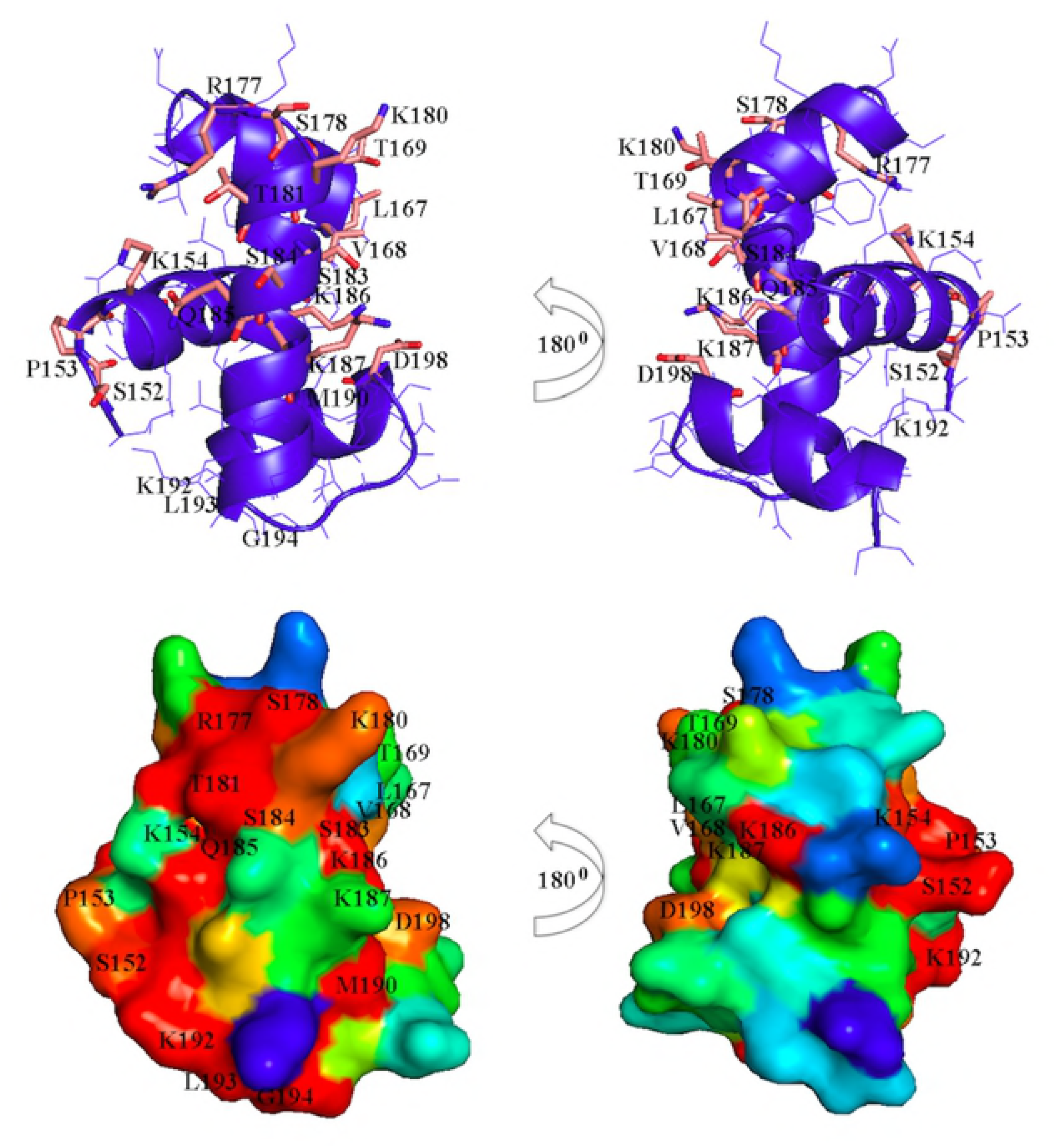
Structure analysis of the RcsB C-terminal DNA-binding domain. Ribbon diagram of the RcsB C-terminal DNA-binding domain from *Erwinia amylovora* (top panel). Conserved residues involve in DNA contacts in known LuxR/FixJ regulators are shown as stick models. RcsB DNA-binding domain: sequence-structure alignment (bottom panel). Surface representation of the RcsB DNA-binding domain was colored by the degree of sequence conservation from red (100% conserved residues) to blue (non-conserved residues). A search for RcsB C-terminal DNA-binding domain homologs was done using the PSI-BLAST server. From the list of 500 sequences against the non-redundant database a random set of 30 sequences with identity from 98% to 40% were chosen. A multiple sequence alignment for visualization of the sequence conservation with respect to the three-dimensional structure was generated.

### SpeG does not bind the LuxR/FixJ family member RcsA

To determine if binding to SpeG is specific for RcsB or if SpeG can bind *in vitro* to other LuxR/FixJ transcriptional regulators that have C-terminal domains similar to RcsB, we heterologously expressed and purified the *E. coli* RcsA transcriptional regulator (an auxiliary partner with RcsB in a heterodimer that interacts with a specific DNA site called the “RcsAB” box [60]) and tested RcsA binding to SpeG by SPR. We found that SpeG does not bind to RcsA in the absence of spermidine (**S4 Fig**), which highlights the binding specificity between RcsB and SpeG. However, we cannot exclude that possibility that SpeG may bind an RcsB-RcsA heterodimer or other LuxR/FixJ-type family members in the presence or absence of spermidine. While RcsA has an HTH motif, its inability to bind SpeG also suggests that other regions of RcsB within its DNA-binding domain besides the HTH motif and/or its oligomeric state might be important for the specificity of the SpeG-RcsB interaction.

## Discussion

We have presented evidence that the metabolic enzyme SpeG regulates transcription from the *rprA* promoter. We also have shown that SpeG binds the DNA binding domain of the transcription factor RcsB. We propose that this interaction interferes with the ability of RcsB to activate transcription from the *rprA* promoter. This represents the first report of a direct link between spermidine metabolism and an envelope stress signal transduction pathway.

### SpeG can inhibit *rprA* transcription through interactions with RcsB

We began this study because we had previously reported that N^ε^-lysine acetylation regulates RcsB activity at the *rprA* promoter [29]. Since deletion of the only known *E. coli* N^ε^-lysine acetyltransferase YfiQ had no obvious effect on the acetylation state of RcsB [29], we screened the known and putative acetyltransferases for regulators of *rprA* transcription and found that SpeG inhibited *rprA* promoter activity: overexpression of SpeG reduced *rprA* promoter activity (**Fig 1A**), while deleting *speG* relieved inhibition of the *rprA* promoter (**Fig 1B**).

Because we did not observe acetylation of RcsB by SpeG (**Fig 2**) and since we did not find that SpeG activity could affect RcsB-dependent *rprA* inhibition (**Fig 3**), we instead investigated the possibility of a physical interaction between RcsB and SpeG. Indeed, SPR analysis showed that SpeG forms a complex with RcsB through the RcsB C-terminal DNA-binding domain (**Fig 4**). We further report that this interaction is specific, as we did not detect binding between SpeG and RcsB’s auxiliary transcription factor RcsA (57) (**S3 and S4 Figs**). These *in vitro* results combined with the *in vivo* analysis support the hypothesis that SpeG and RcsB interact and that the resulting complex impacts RcsB activity at the *rprA* promoter. As *rprA* transcription absolutely requires RcsB, we did not test if SpeG affected *rprA* transcription in an *rcsB* mutant.

### Physiological implications of SpeG-RcsB interactions

It has been estimated that RcsB regulates 5% of the *E. coli* genome, including but not limited to the colanic acid biosynthetic locus, the small RNA *rprA*, and the operon that encodes FlhDC, the master regulator of flagellar biogenesis [61, 62]. The Rcs phosphorelay has also been implicated in regulating biofilm formation and sensitivity to antibiotic-induced peptidoglycan damage [16, 62-64]. Since the SpeG interaction with RcsB regulates activation of *rprA* transcription, SpeG likely influences these other RcsB-regulated phenotypes. In fact, it has been reported that polyamines can induce the glutamate-dependent acid response system [65], which requires RcsB [66]. The outstanding question is why polyamine neutralization and cellular processes regulated by RcsB would be coordinated. We conjecture that SpeG works through members of the RcsB regulon required to initiate proper responses to particular extracellular conditions such as cold shock, heat shock, ethanol, and increased alkalinity, which were shown to influence the spermidine metabolic pathway [67].

## Conclusion

We have shown that SpeG and RcsB can form a complex, suggesting a coordinated response between polyamine metabolism and envelope stress. It is not known why an enzyme involved in spermidine metabolism regulates RcsB, if RcsB affects spermidine biosynthesis, or whether SpeG acts as a general modulator of response regulators. However, it is clear that SpeG inhibits RcsB activity *in vivo* and we propose that it is through a direct interaction between SpeG and the DNA binding domain of RcsB.

## Materials and Methods

### Bacterial strains, bacteriophage, and plasmids

All of the bacterial strains, bacteriophage, and plasmids used in this study are listed in **Table 1**. Derivatives were constructed by generalized transduction with P1kc [68]. P*rprA142*-*lacZ*, a transcriptional fusion of the *rprA* promoter (*PrprA*) to the *lacZ* reporter, was from Dr. Susan Gottesman (National Institutes of Health, Bethesda, MD) [12]. Construction of monolysogens was performed and verified as described previously [69]. Transformations were performed by electroporation or through the use of either transformation buffers 1 and 2 [70] or transformation-and-storage solution [71].

### Culture conditions

For strain construction, cells were grown in lysogeny broth (LB) consisting of 1% (w/v) tryptone, 0.5% (w/v) yeast extract, and 0.5% (w/v) sodium chloride; LB plates contained 1.5% agar. For promoter activity assays, cells were grown in tryptone broth buffered at pH 7 (TB7), which contains 1% (w/v) tryptone buffered at pH 7.0 with potassium phosphate (100 mM). Cell growth was monitored spectrophotometrically (DU640; Beckman Instruments, Fullerton, CA) by determining the absorbance at 600 nm (OD_600_). Chloramphenicol (25 μg/ml) was added to growth media when needed to maintain pCA24n plasmid derivatives. To induce the expression of genes carried on various plasmids, 10 μM isopropyl-β-D-thiogalactopyranoside (IPTG) was added to the growth media.

### β-galactosidase assay

To monitor the promoter activity of P*rprA-lacZ*, biological replicates were grown aerobically at 37°C in TB7 overnight. The overnight cultures were diluted in fresh TB7 to an OD_600_ of 0.05 and grown aerobically with agitation at 250 rpm at 37°C until early stationary phase. At regular intervals, cells were harvested and stored at 4°C in a microtiter plate. β-galactosidase activity was determined quantitatively as described previously (26) using All-in-One β-galactosidase reagent (Pierce Biochemical). Sterile TB7 was used as a negative control on each microtiter plate. Promoter activity was monitored throughout growth and plotted against OD_600_. Each individual experiment included at least three biological replicates. Each of these experiments was performed at least three times. The values represent the means with standard deviations.

### Site-directed mutagenesis

Site-directed mutagenesis of SpeG to pCA24n-speG(Y135A) was conducted in pCA24n-*speG* with the QuikChange Lightning Multi site-directed mutagenesis kit (Agilent Technologies), in accordance with the manufacturer’s instructions by using the mutagenic primers SDMspeGY135A and SDMspeGY135A_as, as listed in **Table 1**.

### RcsB, RcsA and SpeG expression plasmids

Plasmids containing genes from *Escherichia coli* str. K-12 substr. MG1655 included the following: 1) full-length RcsB (NCBI accession code AAC75277, GI: 1788546) in the pCA24n vector from the ASKA collection (chloramphenicol resistant; pCA24n-*rcsB*) [51], 2) the N-terminal receiver domain of RcsB (truncated construct, residues 1-147) in the pMCSG7 vector (ampicillin resistant; pMCSG7-*rcsB*(NTD)), 3) the C-terminal DNA binding domain of RcsB (truncated construct, residues 128-216) in the pMCSG7 vector (ampicillin resistant; pMCSG7-*rcsB*(CTD)), 4) full-length RcsA (NCBI accession code WP_000104001 and GI: CTS77413) in the pMCSG53 vector (ampicillin resistant; pMCSG53-*rcsA*) and 5) full-length SpeG (NCBI accession code NP_416101, GI: 16129542) in the pMCSG7 vector (ampicillin resistant; pMCSG7-*speG*). The full-length RcsB construct (pCA24n-*rcsB*) and its truncated versions ((pMCSG7-*rcsB*(NTD) and pMCSG7-*rcsB*(CTD)) had an uncleavable N-terminal polyhistidine tag, while the RcsA (pMCSG53-*rcsA*) and SpeG (pMCSG7-*speG*) constructs had a cleavable N-terminal polyhistidine tag followed by a TEV protease cleavage site [51]. The genes for the individual RcsB domains were synthesized by Genescript and subcloned into the pMCSG7 vector using ligation independent cloning as described previously [72, 73]. A portion of the linker sequence (comprised of residues 121-149) between the domains was included in each individual domain construct.

### Large-scale protein expression and purification

Expression plasmids containing the desired genes were transformed into kanamycin-resistant BL21(DE3)-magic or KRX/pGro7 (for pMCSG53-*rcsA*) competent cells [74]. pCA24n-*rcsB* transformants were grown in Terrific Broth (TB) in the presence of 34 μg/mL chloramphenicol and 35 μg/mL kanamycin. pMCSG7-*rcsB*(NTD) and pMCSG7-*rcsB*(CTD) transformants were grown in LB in the presence of 400 μg/mL ampicillin and 25 μg/mL kanamycin. The pMCSG53-*rcsA* transformant was grown in M9 L-selenomethionine supplemented media (Medicilon Inc.) in the presence of 400 μg/mL ampicillin, 35 μg/mL kanamycin and 0.1% arabinose. pMCSG7-*speG* transformants were grown in TB supplemented with 100 μg/mL ampicillin and 50 μg/mL kanamycin.

Full-length RcsB protein, its N-terminal and C-terminal domains, and RcsA protein were prepared at the Recombinant Protein Production Core (rPPC) Facility at Northwestern University (Evanston, IL, USA). Transformants containing the RcsB and RcsA plasmids were grown at 37°C in a fermenter until the OD_600_ reached 0.8, whereupon they were induced with 0.6 mM IPTG. The RcsA transformant was also exposed to 0.25% L-rhamnose. The RcsB constructs were expressed at 25°C overnight, whereas RcsA was expressed at 22°C overnight. The next day cells were harvested by centrifugation and resuspended in lysis buffer (1.5 mM magnesium acetate, 1mM calcium chloride, 250 mM sodium chloride, 100 mM ammonium sulfate, 40 mM disodium phosphate, 3.25 mM citric acid, 5% glycerol, 5 mM imidazole, 5 mM beta-mercaptoethanol (BME), 0.08% n-dodecyl-beta-maltoside (DDM), 1 mM phenylmethylsulfonyl fluoride (PMSF), and 20 μM leupeptin) and homogenized. Cells containing the SpeG plasmid were grown at 37°C in a benchtop shaker to an OD_600_ of 0.6, induced with 0.5 mM IPTG, and expressed at 25°C overnight. Cells were harvested and resuspended in lysis buffer (as stated above without PMSF and leupeptin) and sonicated. After sonication, lysates were centrifuged and the supernatant was purified as follows.

The proteins were purified using an ÄKTAxpress™ (GE Healthcare, Piscataway, NJ) high-throughput purification system at 4°C. The crude extract was loaded onto a 5 mL HisTrap FF Ni-NTA column, washed with loading buffer (10 mM Tris HCl pH 8.3, 500 mM sodium chloride and 5 mM BME), washed with loading buffer plus 25 mM imidazole to remove impurities, and eluted with loading buffer plus 500 mM imidazole. The purified proteins were subsequently loaded onto and eluted from a HiLoad™ 26/60 Superdex™ 200 size-exclusion column in loading buffer. The polyhistidine tag of the SpeG protein was removed, as described previously [75]; for all other constructs, the tag remained attached. The final purity of each protein was assayed by SDS-PAGE.

### Enzyme kinetic assays

To test whether SpeG could acetylate RcsB, we performed *in vitro* enzyme kinetics, using a previously described assay and recombinantly expressed and purified proteins [7, 54]. The total volume for each reaction was 50 μL and contained 50 mM Bicine pH 9.0, 0.5 mM acCoA, 1 mM spermidine, 0.96 μM SpeG enzyme, and/or 0.1 mM RcsB full-length protein. All reactions were initiated with 10 μL of SpeG enzyme or enzyme dilution buffer (100 mM Bicine pH 9.0, 100 mM sodium chloride) and were performed in triplicate at 35°C for 20 min. To stop the reactions, 50 μL of a solution containing 100 mM Tris HCl pH 8.0 and 6 M guanidine HCl was added to each reaction. To detect the product of the reaction (CoA), 200 μL of a solution containing 0.2 m*M* 5,5′-Dithiobis(2-nitrobenzoic acid), 100 m*M* Tris HCl pH 8.0, and 1 mM EDTA was added to each reaction and incubated for 10 min at room temperature. The absorbance was then measured at 415 nm on a Biotek ELx808 microplate reader.

### Surface plasmon resonance (SPR) analysis of binding interactions between SpeG and RcsB in absence or presence of spermidine

Binding interactions of *E. coli* SpeG to full-length *E. coli* RcsB or individual RcsB domains in the absence of spermidine were measured using a Reichert SR7500DC (Reichert Technologies, Buffalo, NY) dual channel spectrometer at the Keck Biophysics Facility at Northwestern University (Evanston IL, USA). Prior to immobilizing SpeG onto a carboxymethyl dextran hydrogel surface gold sensor chip (Reichert Technologies, Buffalo, NY), the surface of the chip containing COO^-^ groups were activated with a mixture of N-hydroxysuccinimide (NHS) and 1-ethyl-3-(3-dimethylaminopropyl)carbodiimide hydrochloride (EDC) to create amine reactive esters. SpeG protein (46 μM) in solution containing 10 mM HEPES at pH 8.3 and 100 mM sodium chloride was then immobilized onto the chip and covalently coupled with the surface NHS esters at a flow rate of 40 μL/min at room temperature. To achieve saturation, two sequential injections of SpeG for 3 min followed by 1.5 min of dissociation were performed. To block formation of residual NHS esters, an ethanolamine solution was injected over the chip. To remove weakly bound SpeG molecules, the chip was washed with running buffer containing 10 mM HEPES at pH 8.3 and 100 mM sodium chloride. The instrument was cooled and all SPR measurements were carried out at 4°C. All protein solutions were prepared in running buffer. 160 μL of RcsB full-length (10, 21, 42, 53, 63 and 74 μM), RcsB C-terminal domain (45, 67, 91, 114, 136, and 159 μM), or RcsB N-terminal domain (59, 118, and 176 μM) were injected sequentially over the SpeG-chip with a flow rate of 40 μL min^-1^ for 30 sec followed by a 1.5 min rinse and a 1 min dissociation. After each binding cycle, SpeG surfaces were regenerated by injecting 0.5 M sodium chloride for 45 sec at a flow rate of 30 μL min^-1^ and washed with running buffer. All analyte injections were performed in duplicate. For each measurement, a background response recorded in the reference cell was subtracted as well as the response from a blank injection with the running buffer.

To investigate how spermidine affects binding interactions of SpeG to RcsB and its individual domains, we used a Reichert4SR (Reichert Technologies, Buffalo, NY) four-channel SPR system at the Keck Biophysics Facility. SpeG was immobilized at a concentration of 46 μM onto cells 3 and 4 using the amine coupling procedure described above. Cells 1 and 2 were used as reference cells. All measurements were performed at 4°C. A solution containing 0.5 mM spermidine in the running buffer was flowed over the surface of the immobilized SpeG at 40 μL min-1 for 30 sec followed by a 1.5 min rinse and a 1 min dissociation. The chip was then washed with running buffer until the SPR signal reached a stable value. 160 μL of RcsB full-length (10, 21, 42, 53, 63 and 74 μM), RcsB C-terminal domain (23, 45, 91, 114, and 136 μM) or RcsB N-terminal domain (59, 118, 177, 236 and 295 μM) were injected sequentially over the SpeG-chip, as described above, to monitor binding of RcsB constructs to SpeG in the presence of spermidine. After each binding cycle of RcsB full-length and RcsB C-terminal domain, SpeG surfaces were regenerated with an injection of 0.5 M sodium chloride for 1.5 min at a flow rate of 30 μL min-1 and washed with the running buffer. Regeneration of the chip surface after injections of RcsB N-terminal domain was not required because the protein dissociated on its own. A background response for each run and the response from a blank injection were subtracted. With the exception of the RcsB N-terminal domain in the absence of spermidine, duplicate measurements were collected for each concentration of each protein. Data processing and kinetic analyses for all experiments were performed using TraceDrawer Data Analysis software (Reichert Technologies, Buffalo, NY).

### SPR analysis of binding interactions between SpeG and the transcription factor RcsA

To examine binding interactions between SpeG and RcsB’s auxiliary partner RcsA from *E. coli*, we used a four-channel SPR system at the Keck Biophysics Facility following the amine coupling protocol, as described above. SpeG protein at a concentration of 46 μM in 10 mM HEPES buffer at pH 8.3 containing 100 mM sodium chloride was immobilized onto the chip. 160 μL of RcsA protein solution in running buffer (21, 42, 64 and 85 μM) was injected consecutively over the SpeG-chip followed by regeneration and washing, as described above. A background response and response from a blank injection that contained running buffer were subtracted from each sensorgram to determine the actual binding response. Data were processed using TraceDrawer software.

### Linear Regression Analysis

To determine whether experimental results were statistically significant, a linear regression was performed, comparing all experimental groups with their respective vector controls. All of the regressions used were set up as follows: the calculated *rprA* promoter activity was the response variable, the overexpressed plasmids or mutant were the explanatory variable, and time was a random effect. OD was not included as an effect on activity as it is already used in the calculation of activity. Time as a random effect was chosen based on the question asked: Accounting for the effects of time on activity does the experimental group in question significantly affect overall *rprA* promoter activity? The significance threshold was set at 0.05. The open source program R (version 3.3.2) and packages “lmerTest”, “ggplot2”, and “moments” were used to visualize and analyze the data (76,77,78,79).

## Acknowledgements

We would like to thank the Keck Biophysics Facility at Northwestern University (Evanston, IL) for assistance with SPR data collection, Roberto Limeira and Cara Joyce (Loyola University Chicago) for the linear regression analyses of the *in vivo* data, Sarah Cook, David Christensen, and Bozena Zemaitaitis (Loyola University Chicago) for helping to complete and repeat the acetyltransferase overexpression experiments, and the members of the Visick and Wolfe labs (Loyola University Chicago) for important scientific discussions pertaining to this study.

## Supporting information

**S1 Fig. The effect of spermidine on rprA in the absence of SpeE.**

The *speE* mutant was transformed with either the VC or pSpeG and grown in TB7 supplemented with 50 μM IPTG and 0, 1.5, 2.5, or 5 mM spermidine. Growth and *rprA* promoter activity was measured over time. Each data point is an average of duplicate biological replicates and standard deviations.

**S2 Fig. Affinity analysis of RcsB binding to SpeG.**

(A) The maximum responses in the SPR sensograms for the first dilution series of RcsB C-terminal domain in the absence of spermidine are plotted against the analyte concentration. (B and C) The SPR sensograms for dilution series of RcsB full-length and its C-terminal domain after exposure to spermidine. The RcsB full-length or RcsB C-terminal domain protein was injected in five dilution series with the following concentrations: 21, 42, 53, 63 and 74 μM (B) or 23, 45, 91, 114, and 136 μM (C). The fitted data are shown in black.

**S3 Fig. Phylogenetic tree of LuxR/FixJ DNA-binding domain of transcriptional regulators.**

Phylogenetic tree was created in ClustalW2 server (http://www.ebi.ac.uk/Tools/msa/clustalw2). A list of 58 representatives of the conserved LuxR/FixJ DNA-binding domains was generated in NCBI server http://www.ncbi.nlm.nih.gov/Structure/cdd) and includes DNA-binding domains of following transcriptional factors: RcsB from *Escherichia coli* (RcsB-Es_co) [GI:353570681], RcsB from *Erwinia amylovora* (RcsB-Er_am) [GI:33357861], YjjQ from *E. coli* (YjjQ-Es_co) [GI:83288197], BglJ from *E. coli* (BglJ-Es_co) [GI:3915634], YahA from *E. coli* (YahA-Es_co) [GI:2506596], YuaB from *E. coli* (YuaB-Es_co) [GI:81783897], DctR from *E. coli* (DctR-Es_co) [GI:57012697], RcsA from *E. coli* (RcsA-Es_co) [GI:60393000], EntR from *Citrobacter freundii* (EntR-Ci_fr) [GI:6015049], FimW from *Salmonella enterica* (FimW-Sa_en) [GI:585140], LuxR from *Bacteroides thetaiotoamicron* (LuxR-Ba_th) [GI:171849138], YgeK from *E. coli* (YgeK-Es_co) [GI:20140955], UhpA from *E. coli* (UhpA-Es_co) [GI:84029412], UvrY from *E. coli* (UvrY-Es_co) [GI:83288180], PA0034 from *Pseudomonas aeruginosa* (PA0034-Ps_ae) [GI:13959718], BvgA from *Bordetella pertussis* (BvgA-Bo_pe) [GI:61219948], FimZ from *E. coli* (FimZ-Es_co) [GI:84028128], EvgA from *E. coli* (EvgA-Es_co) [GI:82581667], FixJ from *Sinorhizobium meliloti* (FixJ-Si_me) [GI:159163516], StyR from *P. fluorescens* (StyR-Ps_fl) [GI:78100993], NodW from *Bradyrhizobium diazoefficiens* (NodW-Br_di) [GI:128495], Ycf29 from *Porphyra purpurea* (Ycf29-Po_pu) [GI:1723332], Ycf29 from *Cyanophora paradoxa* (Ycf29-Cy_pa) [GI:1351750], NarL from *E. coli* (NarL-Es_co) [GI:24158735], NarP from *E. coli* (NarP-Es_co) [GI:400374], GerE from *Bacillus subtilis* (GerE-Ba_su) [GI:13786948], VraR from *Staphylococcus aureus* (VraR-St_au) [GI:166007196], LiaR from *B. subtilis* (LiaR-Ba_su) [GI:68051995], DegU from *B. subtilis* (DegU-Ba_su) [GI:118438], YxjL from *B. subtilis* (YxjL-Ba_su) [GI:20141933], YhjB from *E. coli* (YhjB-Es_co) [GI:586682], CsgD from *E. coli* (CsgD-Es_co) [GI:1706166], MoaR from *Enterobacter aerogenes* (MoaR-En_ae) [GI:1709068], MalT from *E. coli* (MalT-Es_co) [GI:189028606], SgaR from *Hyphomicrobium methylovorum* (SgaR-Hy_me) [GI:6094276], Rv08090c from *Mycobacterium tuberculosis* (Rv08090c-My_tu) [GI:6137301], AgmR from *P. aeruginosa* (AgmR-Ps_ae) [GI:121420], AlkS from *P. oleovorans* (AlkS-Ps_ol) [GI:6226550], ComA from *B. subtilis* (ComA-Ba_su) [GI:116903], YdfI from *B. subtilis* (YdfI-Ba_su) [GI:68566110], ExeN from *Aeromonas salmonicida* (ExeN-Ae_sa) [GI:1175862], LuxR from *Aliivibrio fischeri* (LuxR-Al_fi) [GI:462556], VanR from *Vibrio anguillarum* (VanR-Vi_an) [GI:9297072], SolR from *Ralstonia solanacearum* (SolR-Ra_so) [GI:9297032], AhyR from *Aeromonas hydrophila* (AhyR-Ae_hy) [GI:61218504], LasR from *P. aeruginosa* (LasR-Ps_ae) [GI:125980], Y4HQ from *Sinorhizobium fredii* (Y4HQ-Si_fr) [GI:2495427], SdiA from *E. coli* (SdiA-Es_co) [GI:2506570], PhzR from *P. fluorescens* (PhzR-Ps_fl) [GI:2495423], CarR from *Pectobacterium carotovorum* (CarR-Pe_ca) [GI:2495418], YenR from *Yersinia enterocolitica* (YenR-Ye_en) [GI:1723596], RhiR from *Rhizobium leguminosarum* (RhiR-Rh_le) [GI:417645], TraR from *S. fredii* (TraR-Si_fr) [GI:158429605], MoxX from *Paracoccus denitrifican* (MoxX-Pa_de) [GI:266552], BrpA from *Streptomyces hygroscopicus* (BrpA-St_hy) [GI:231653], RaiR from *Rhizobium etli* (RaiR-Rh_et) [GI:9297035], TraR from *Agrobacterium tumefaciens* (TraR-Ag_tu) GI:23200109 and TraJ from *E. coli* (TraJ-Es_co) [GI:464931].

**S4 Fig. SPR analysis of SpeG and RcsA interaction.**

The SPR sensograms of SpeG and transcription regulator RcsA. The RcsA protein was injected in four dilution series. Duplicate measurements for each concentration indicated above SPR sensograms were performed.

